# Oxytocin regulates TN-GnRH3 circuit maturation and mate preference through C1q-dependent synaptic mechanisms

**DOI:** 10.64898/2026.05.17.725056

**Authors:** Maya Hasegawa, Moyu Oshita, Kiyoshi Naruse, Kjetil Hodne, Finn-Arne Weltzien, Shin-ichi Higashijima, Taiju Amano, Masabumi Minami, Hideaki Takeuchi, Shinichi Nakagawa, Saori Yokoi

## Abstract

Oxytocin is a key regulator of social behavior, yet how it shapes the neural circuits underlying these behaviors remains unclear. Here, we show that oxytocin signaling is required for the proper maturation of terminal nerve gonadotropin-releasing hormone 3 (TN-GnRH3) circuits that regulate female mate preference in medaka. Disruption of oxytocin signaling reduced expression of the complement component C1q, altered microglial properties, and increased TN-GnRH3 innervation in the optic tectum and dorsolateral telencephalon, suggesting impaired synaptic refinement. In addition, TN-GnRH3 neurons in mutants exhibited disrupted clustering and failed to increase firing frequency in response to visual stimulation, indicating functional deficits.

Furthermore, *c1qb* mutants phenocopied the behavioral and neuronal abnormalities observed in oxytocin signaling–deficient mutants.

Together, our findings suggest that oxytocin signaling links microglia-associated synaptic refinement to neural circuit maturation, thereby revealing a previously unrecognized role for oxytocin in shaping social behavior.

## Introduction

Oxytocin is a key neuromodulator of social behavior across vertebrates, including social recognition, pair bonding, and social affiliation^1–3^. Despite extensive studies on its role in modulating neuronal activity and behavioral outputs, how oxytocin signaling contributes to the establishment of neural circuits underlying social decision-making remains poorly understood.

Recent studies have suggested that oxytocin can influence not only neuronal function but also non-neuronal cell types. In particular, oxytocin has been reported to modulate microglial activity in the context of inflammation and stress responses, often exerting anti-inflammatory effects in the brain^4,5^. However, whether oxytocin signaling regulates microglial functions involved in neural circuit development remains unclear.

Microglia play a central role in the refinement of neural circuits during development through synaptic pruning^6^. This process is mediated in part by the complement system, in which molecules such as C1q tag synapses for elimination by microglia, thereby shaping the maturation and functional properties of neural networks^7,8^. Disruption of microglia-dependent synaptic refinement has been linked to altered circuit function and behavioral abnormalities^9,10^, yet the upstream signals that regulate this process in the context of social behavior remain largely unknown.

The teleost fish medaka (*Oryzias latipes*) provides a tractable model to investigate the neural basis of visually guided mate preference^11–14^. Female medaka preferentially select visually familiarized males as mating partners, a process that is mediated by increased activity of terminal nerve GnRH3 (TN-GnRH3) neurons upon visual exposure to a male. Perturbation of TN-GnRH3 neuron development leads to defects in mate preference, suggesting that proper maturation of TN-GnRH3 circuits is essential for normal mate preference^13^. Notably, oxytocin signaling–deficient medaka lose their preference for familiarized males, indicating a critical role for oxytocin signaling in mate preference^14^. Our previous transcriptomic analyses suggested that the expression of genes encoding C1q components (*c1qa*, *c1qb*, and *c1qc*) was reduced under conditions of disrupted oxytocin signaling^14^, suggesting that oxytocin influences the maturation of TN-GnRH3 circuits through complement-mediated synaptic refinement.

Here, we investigated whether oxytocin signaling regulates the maturation of TN-GnRH3 circuits through microglia-associated synaptic refinement. We show that loss of oxytocin signaling leads to alterations in microglial morphology and reduced expression of complement genes in the forebrain and midbrain. These changes are accompanied by abnormal TN-GnRH3 neuronal structure and projection patterns, as well as impaired activity responses to male stimuli. Furthermore, disruption of the complement component C1qb phenocopies the behavioral and neuronal defects observed in both oxytocin and oxytocin receptor mutants. Together, our findings suggest that oxytocin signaling contributes to microglia-mediated synaptic refinement underlying TN-GnRH3 circuit function and mate preference.

## Results

### C1q is required for mate preference and reduced in oxytocin signaling–deficient medaka

Our previous bulk RNA sequencing analyses of brain tissues from oxytocin and oxytocin receptor mutant medaka revealed reduced expression of all complement component C1q genes, including *c1qa, c1qb,* and *c1qc*^14^. As C1q plays a central role in complement-mediated synaptic refinement, these findings raised the possibility that altered C1q signaling contributes to the behavioral defects observed in oxytocin signaling–deficient medaka.

To test whether disruption of the C1q pathway underlies the impaired mate preference in these mutants, we generated medaka lacking c1q function. As C1q is composed of a heterotrimer formed by C1qa, C1qb, and C1qc subunits^15^, we focused on *c1qb* and established mutant lines for further analysis. We generated knockout medaka lines using CRISPR/Cas9, leading to a 5-bp insertion in the second exon. This mutation resulted in a frameshift and subsequent loss of the C1q domain, suggesting a loss-of-function mutation in these medaka (Fig. 1a).

**Fig. 1.**
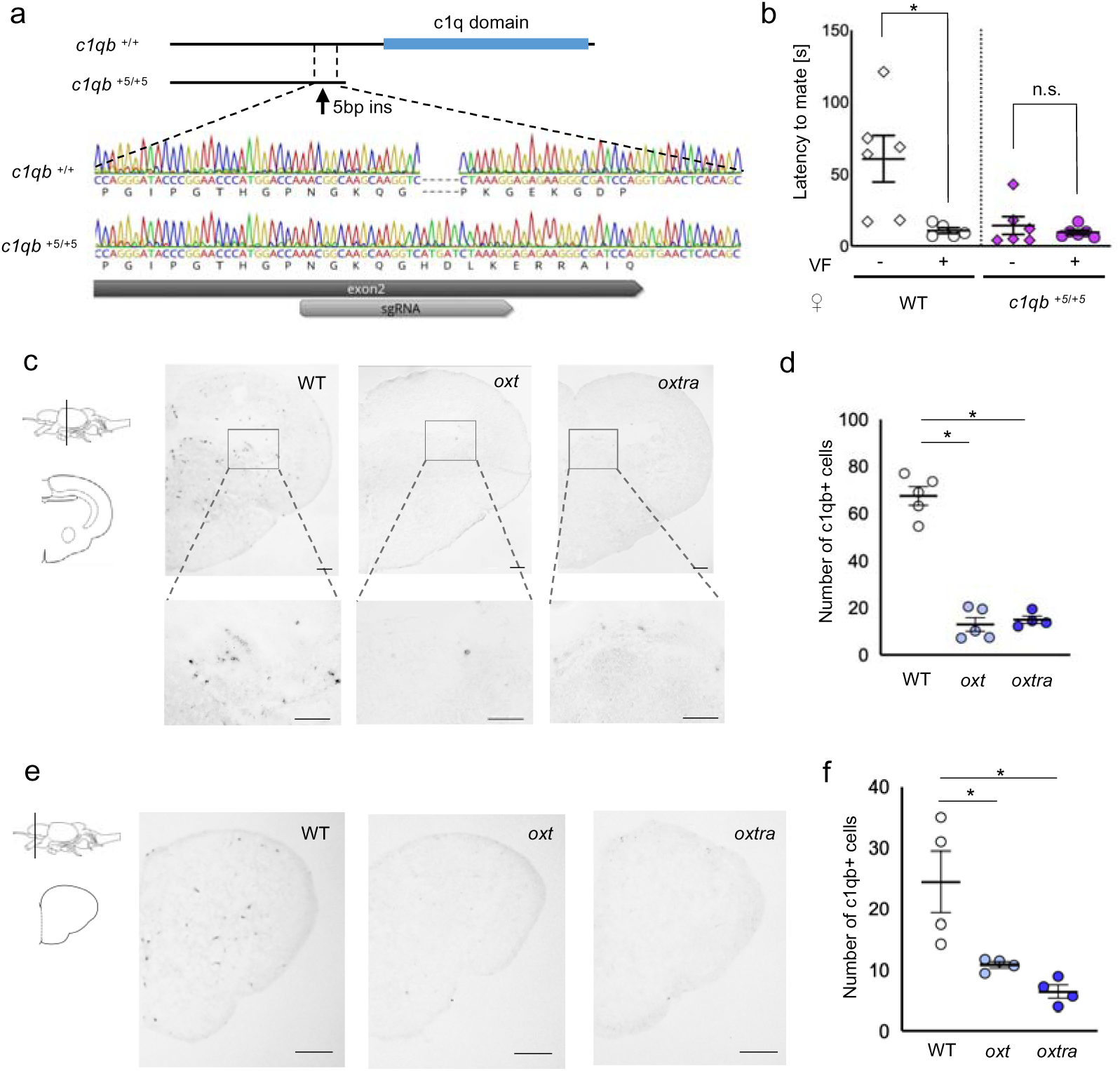
Reduced *c1qb* expression in oxytocin signaling–deficient medaka and its role in mate preference. a. Outline of *c1qb* mutant (*c1qb ^+5/+5^*). In this mutant, the c1qb gene is likely truncated, resulting in the loss of a c1q domain. A 5 bp insertion in exon 2 of the mutant gene lead to the incorporation of ten residues with an altered sequence. In total, 40 residues were added to the altered sequence, followed by a stop codon in this mutant. b. Mating preference of *c1qb* mutant females. While visual familiarization (VF) increased the receptivity of WT females toward familiarized males, *c1qb* mutant females showed high receptivity even toward unfamiliar males. Mean ± SEM, n = 6, 5, 6, 6. Mann–Whitney U test: *P < 0.05; n.s., not significant. c. Representative micrographs showing *c1qb* expression in the midbrain of WT, *oxt* mutant, and *oxtra* mutant females. The position of the coronal section is indicated by the line. Higher-magnification views of the boxed regions are shown below. Scale bars represent 70 μm. d. The number of *c1qb*-positive cells in the midbrain of WT, *oxt* mutant, and *oxtra* mutant females. Mean ± SEM, n = 5, 5, 4. Kruskal–Wallis test followed by Steel’s post hoc test. *P < 0.05. e. Representative micrographs showing *c1qb* expression in the forebrain of WT, *oxt* mutant, and *oxtra* mutant females. The position of the coronal section is indicated by the line. Scale bars represent 100 μm. f. The number of *c1qb*-positive cells in the forebrain of WT, *oxt* mutant, and *oxtra* mutant females. Mean ± SEM, n = 4, 4, 4. Kruskal–Wallis test followed by Steel’s post hoc test. *P < 0.05.

To examine the role of C1q in mate preference, we performed the behavioral assay using female *c1qb* mutants. In wild-type (WT) females, visual familiarization (VF), in which a male and a female were allowed visual contact through a transparent partition before testing, significantly decreased the latency to mating compared to the no-VF condition. The latency to mating was defined as the interval between the first male courtship behavior and the first mating event. This reduction in latency was correlated with increased female receptivity, indicating a preference for visually familiarized males^13,14^. In contrast, *c1qb* mutant females exhibited a short latency to mating even in the absence of VF and displayed high receptivity toward both familiarized and unfamiliar males (Fig. 1b). These results indicate that *c1qb* mutants lose mate preference and instead show indiscriminate mating behavior. Notably, this phenotype closely resembles that observed in oxytocin signaling–deficient females^14^, suggesting that disruption of the C1q pathway phenocopies the behavioral defects caused by impaired oxytocin signaling.

Because *c1qb* mutants phenocopied the behavioral defects observed in oxytocin signaling–deficient medaka, we next examined *c1qb* expression at the cellular level using *in situ* hybridization in oxytocin (*oxt*) and oxytocin receptor (*oxtra*) mutant medaka lines^14^. TN-GnRH3 neurons are key regulators of mate preference behavior^13^, and we therefore asked whether *c1qb* expression is altered in regions associated with TN-GnRH3 circuits. TN-GnRH3 neurons are located in the forebrain and project to the midbrain and the other forebrain regions^16,17^, and we focused on these regions to assess changes in *c1qb* expression. In WT fish, *c1qb*-expressing cells were broadly distributed in both the forebrain and midbrain (Figs. 1c, e). In contrast, the number of *c1qb*-positive cells was significantly reduced in both regions in *oxt* and *oxtra* mutant brains compared to WT (Figs. 1c-f). These results suggest that oxytocin signaling is required for normal levels of *c1qb* expression.

### Microglial morphology is altered in oxytocin signaling–deficient and *c1qb* mutant medaka

As C1q has been reported to be expressed in microglia in other vertebrates^18^, we first examined whether this is also the case in medaka. We performed combined *in situ* hybridization for *c1qb* transcripts and immunostaining for the microglial marker Iba1. We observed *c1qb* signals in Iba1-positive cells, indicating that C1q is expressed in microglia in medaka (Fig. S1).

We next examined whether microglial properties are altered in *oxt*, *oxtra*, and *c1qb* mutant medaka. Consistent with the reduced *c1qb* expression observed in both the forebrain and midbrain of *oxt* and *oxtra* mutants, we analyzed microglia in these regions. Immunostaining for Iba1 revealed no significant difference in the number of microglia between WT and mutant brains across all three mutant lines in either region (Figs. 2a-b, d-e). In contrast, higher-magnification images suggested morphological differences between WT and mutant microglia, with WT microglia appearing relatively amoeboid-like and mutant microglia exhibiting more ramified morphologies (Fig. 2a). Quantitative analysis showed that the average cell area of microglia was significantly larger in WT compared to mutants in both the forebrain and midbrain (Figs. 2c, f). This difference is consistent with morphological differences between WT and mutant microglia, with mutant microglia appearing more ramified than those in WT brains. As microglial morphology is associated with their functional state^19^, these results suggest that microglial functional states may be altered in these mutants. We did not detect expression of the oxytocin receptor in microglia (Fig. S2), suggesting that oxytocin signaling may influence microglial properties indirectly.

**Fig. 2.**
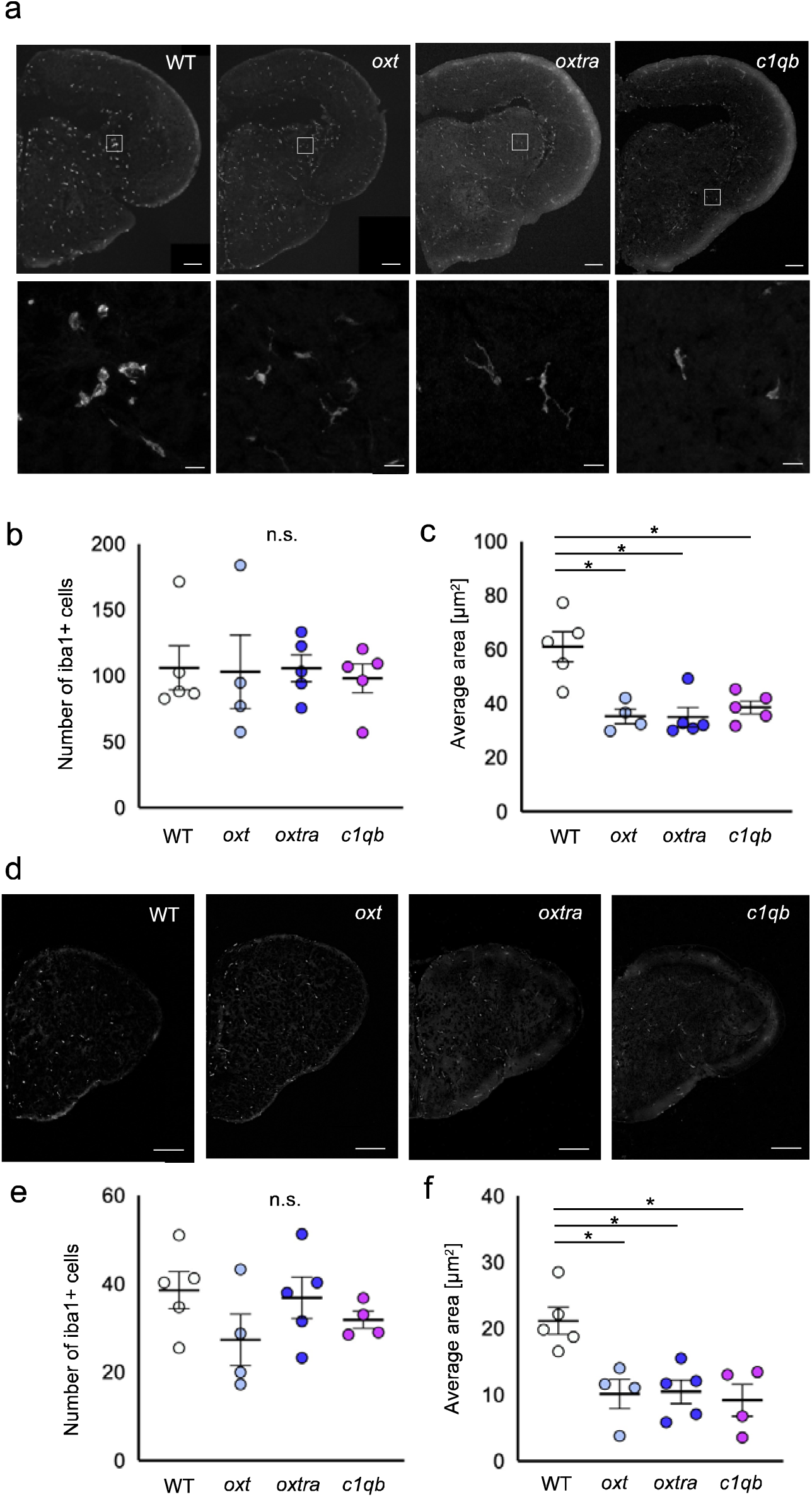
Microglial morphology, but not cell number, is altered in mutants. a. Representative micrographs showing Iba1 expression in the midbrain of WT, *oxt* mutant, *oxtra* mutant, and *c1qb* mutant females. The position of the coronal section corresponds to that shown in Fig. 1c (midbrain). Higher-magnification views of the boxed regions are shown below. Scale bars represent 100 μm in low-magnification images and 10 μm in higher-magnification images. b. The number of Iba1-positive cells in the midbrain of WT, *oxt* mutant, *oxtra* mutant, and *c1qb* mutant females. Mean ± SEM, n = 5, 4, 5, 5. Kruskal–Wallis test. n.s., not significant. c. The average area of Iba1-positive cells in the midbrain of WT, *oxt* mutant, *oxtra* mutant, and *c1qb* mutant. Mean ± SEM, n = 5, 4, 5, 5. Kruskal–Wallis test followed by Steel’s post hoc test. *P < 0.05. d. Representative micrographs showing Iba1 expression in the midbrain of WT, *oxt* mutant, *oxtra* mutant, and *c1qb* mutant females. The position of the coronal section corresponds to that shown in Fig. 1e (forebrain). Scale bars represent 100 μm. e. The number of Iba1-positive cells in the forebrain of WT, *oxt* mutant, *oxtra* mutant, and *c1qb* mutant females. Mean ± SEM, n = 5, 4, 5, 4. Kruskal–Wallis test. n.s., not significant. f. The average area of Iba1-positive cells in the forebrain of WT, *oxt* mutant, *oxtra* mutant, and *c1qb* mutant. Mean ± SEM, n = 5, 4, 5, 4. Kruskal–Wallis test followed by Steel’s post hoc test. *P < 0.05.

### TN-GnRH3 innervation is increased in mutant medaka

Previous studies have shown that terminal nerve GnRH3 (TN-GnRH3) neurons play a central role in regulating mate preference behavior^13^. Based on our findings that microglial morphology is altered in these mutants, we hypothesized that abnormal microglial function may affect the development of TN-GnRH3 neural circuits. TN-GnRH3 neurons project broadly throughout the brain, including to the optic tectum, where they are thought to modulate visual processing and integrate sensory and reproductive signals to regulate behavior^20,21^. To visualize TN-GnRH3 neurons, we generated transgenic medaka expressing GFP under the control of the gnrh3 promoter^16^. Using this line, we examined TN-GnRH3 innervation in the optic tectum (Fig. 3a). Quantitative analysis revealed that TN-GnRH3 innervation in the optic tectum was significantly increased in *oxt*, *oxtra*, and *c1qb* mutant females compared to WT (Fig. 3b). Similar increases were also observed in the dorsal telencephalon (Dl), a higher-order sensory processing region in the forebrain, including visual processing^22,23^ (Figs. 3c, d). These results suggest that TN-GnRH3 projections are increased in these mutants.

**Fig. 3.**
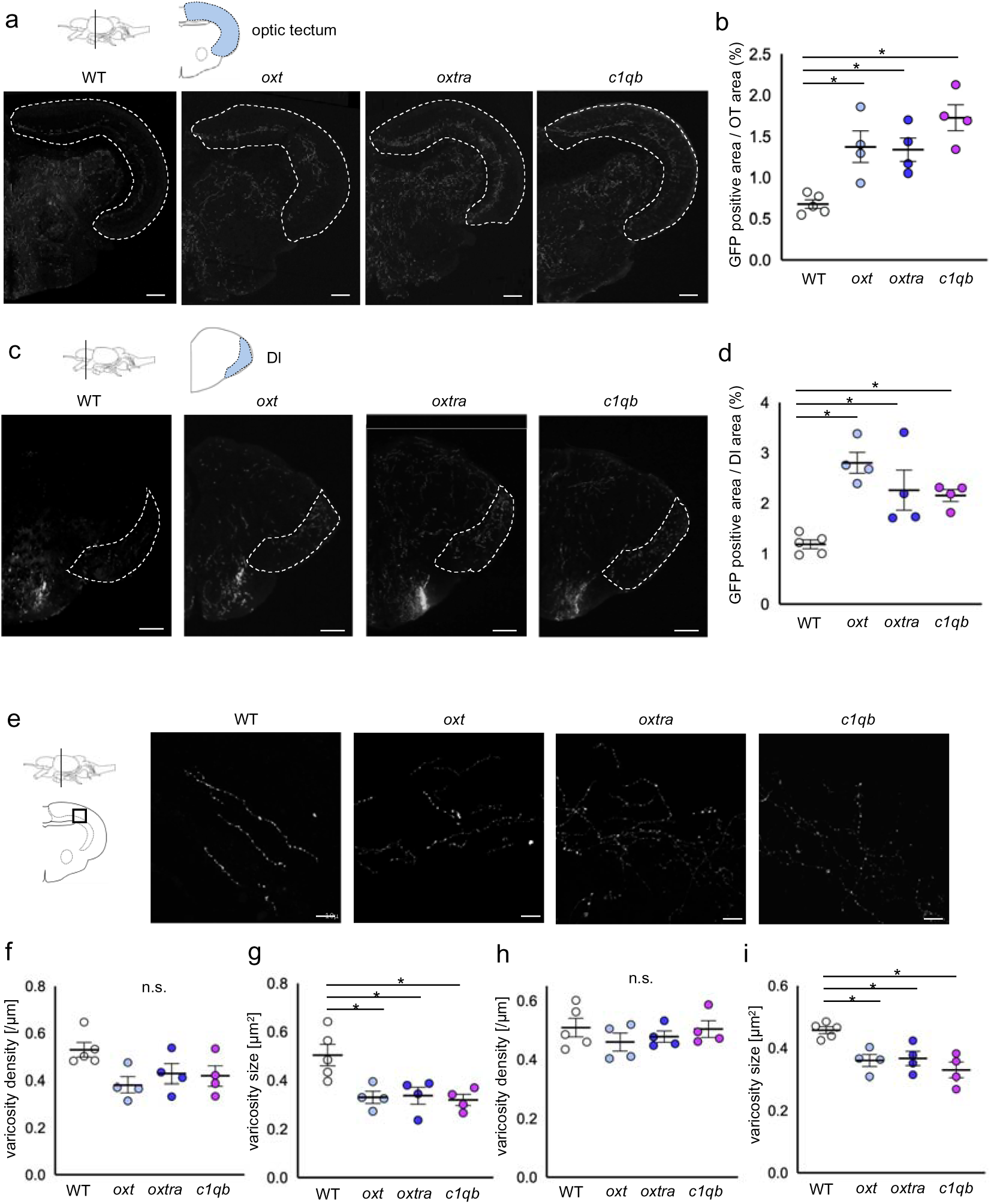
Excessive innervation and reduced varicosity size of TN-GnRH3 neurons in mutants. a. Representative micrographs of gnrh3:GFP fluorescence in the midbrain of WT, *oxt* mutant, *oxtra* mutant, and *c1qb* mutant females. The optic tectum is outlined by a dashed line. Increased innervation in the optic tectum region appears to be higher in mutant medaka compared to WT. The position of the coronal section is indicated by the line. Scale bars represent 100 μm. b. The relative area of GFP-positive signals within the optic tectum (OT) in WT, *oxt* mutant, *oxtra* mutant, and *c1qb* mutant females. Mean ± SEM, n = 5, 4, 4, 4. Kruskal–Wallis test followed by Steel’s post hoc test. *P < 0.05. c. Representative micrographs of gnrh3:GFP fluorescence in the forebrain of WT, *oxt* mutant, *oxtra* mutant, and *c1qb* mutant females. The dorsolateral telencephalon (Dl) is outlined by a dashed line. Increased innervation in the Dl region appears to be higher in mutant medaka compared to WT. The position of the coronal section is indicated by the line. Scale bars represent 100 μm. d. The relative area of GFP-positive signals within Dl in WT, *oxt* mutant, *oxtra* mutant, and *c1qb* mutant females. Mean ± SEM, n = 5, 4, 4, 4. Kruskal–Wallis test followed by Steel’s post hoc test. *P < 0.05. e. Higher-magnification views of the boxed regions, highlighting axonal varicosities of gnrh3:GFP-positive neurons in WT, *oxt* mutant, *oxtra* mutant, and *c1qb* mutant females. Varicosities appear larger in WT and smaller in all three mutant lines. Scale bars represent 10 μm. f. The average density of axonal varicosities in the optic tectum of WT, *oxt* mutant, *oxtra* mutant, and *c1qb* mutant females. Mean ± SEM, n = 5, 4, 4, 4. Kruskal–Wallis test. n.s., not significant. g. The average area of axonal varicosities in the optic tectum of WT, *oxt* mutant, *oxtra* mutant, and *c1qb* mutant females. Mean ± SEM, n = 5, 4, 4, 4. Kruskal–Wallis test followed by Steel’s post hoc test. *P < 0.05. h. Quantification of the average density of axonal varicosities in Dl of WT, *oxt* mutant, *oxtra* mutant, and *c1qb* mutant females. Mean ± SEM, n = 5, 4, 4, 4. Kruskal–Wallis test. n.s., not significant.i. Quantification of the average area of axonal varicosities in Dl of WT, *oxt* mutant, *oxtra* mutant, and *c1qb* mutant females. Mean ± SEM, n = 5, 4, 4, 4. Kruskal–Wallis test followed by Steel’s post hoc test. *P < 0.05.

### Reduced varicosity size in mutant medaka suggests impaired synaptic maturation

To further assess synaptic properties of TN-GnRH3 projections, we attempted to visualize synapses using antibodies against synaptic markers, including synaptophysin and PSD95; however, these approaches did not yield reliable signals in our preparation. We therefore focused on axonal varicosities as an alternative structural proxy for presynaptic sites. Varicosities contain synaptic vesicles and neurotransmitters and can function as *en passant* synapses, representing presynaptic structures^24^. High-magnification analysis of TN-GnRH3 projections in the optic tectum revealed that neurons in mutant fish exhibited smaller varicosities compared to WT (Fig. 3e). Quantitative analysis showed that while the density of varicosities was not significantly different between groups (Fig. 3f), the average size of varicosities was significantly reduced in all three mutant lines (Fig. 3g). Similar results were also observed in Dl, where increased TN-GnRH3 innervation was detected (Figs. 3h, i). These findings suggest that presynaptic structural properties of TN-GnRH3 projections are altered in these mutants, consistent with altered presynaptic structure that may reflect impaired synaptic maturation.

In addition to alterations in axonal innervation, we next examined whether TN-GnRH3 neuronal soma are also affected in these mutants. TN-GnRH3 neurons are known to form a compact cluster and are characterized by relatively large cell bodies^20,25^. We therefore compared these features between WT and mutant fish. In all three mutant lines, TN-GnRH3 neuronal clusters appeared more dispersed compared to WT (Fig. 4a). Quantification of cluster compactness, based on the distance between the farthest cell bodies, confirmed that clusters were more dispersed in mutants (Fig. 4b). In addition, the size of neuronal nuclei tended to be smaller in mutants (Fig. 4c). By contrast, the number of TN-GnRH3 neuronal somata was not significantly different between WT and mutants (Fig. S3). These results suggest that the morphological properties of TN-GnRH3 neurons are altered in these mutants, which may contribute to altered neuronal function. These alterations in TN-GnRH3 neuronal morphology were observed in regions corresponding to those showing reduced *c1qb* expression and altered microglial morphology (Figs. 1e, 2d; see Fig. S4 for section levels containing TN-GnRH3 neuronal somata).

**Fig. 4.**
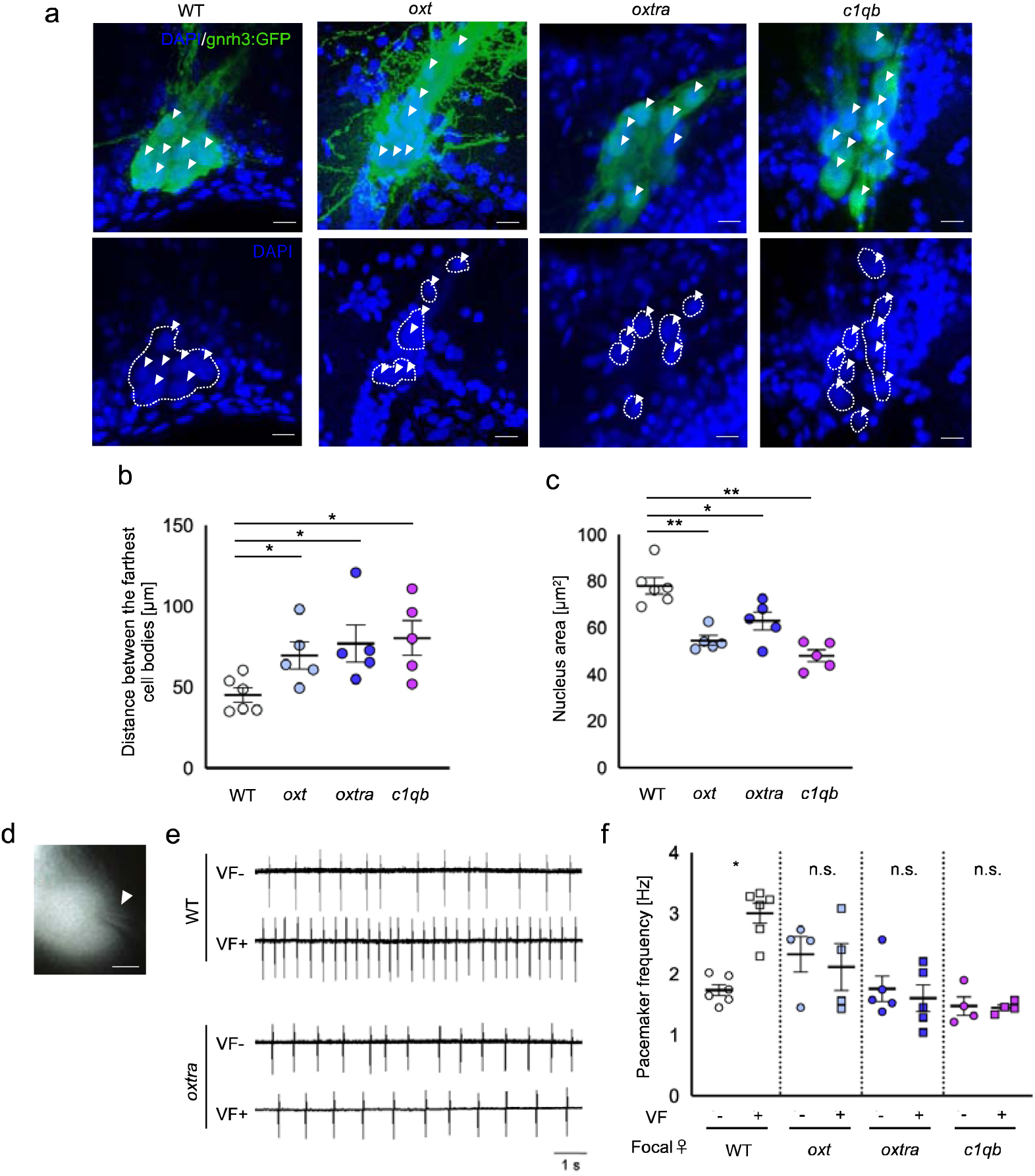
Disrupted clustering and impaired activity of TN-GnRH3 neurons in mutants. a. Representative micrographs of TN-GnRH3 neuronal somata in the forebrain of WT, *oxt* mutant, *oxtra* mutant, and *c1qb* mutant females. Top, gnrh3:GFP (green) merged with DAPI (blue); bottom, DAPI channel of the same region. Arrowheads indicate TN-GnRH3 neuronal cell bodies. Dashed outlines indicate the positions of nuclei. Neuronal clusters appear less compact in mutants, and nuclei tend to be smaller compared to WT. Scale bars represent 10 μm. b. Quantification of the distance between the farthest cell bodies of TN-GnRH3 in WT, *oxt* mutant, *oxtra* mutant, and *c1qb* mutant females. An increase in this distance reflects a less compact (more dispersed) neuronal cluster. Mean ± SEM, n = 6, 5, 5, 5. Kruskal–Wallis test followed by Steel’s post hoc test. *P < 0.05. c. Average nucleus area of TN-GnRH3 in WT, *oxt* mutant, *oxtra* mutant, and *c1qb* mutant females. Mean ± SEM, n = 6, 5, 5, 5. Kruskal–Wallis test followed by Steel’s post hoc test. **P < 0.01, *P < 0.05. d. Representative image of a GFP-labeled TN-GnRH3 neuron during electrophysiological recording. Arrowhead indicates the recording electrode. Scale bars represent 5 μm. e. Representative loose cell-attached recordings from TN-GnRH3 neurons in WT and *oxtra* mutant females with or without visual familiarization (VF) to a male. f. Quantification of pacemaker firing frequency of TN-GnRH3 neurons in focal females under visual familiarization (VF- and VF+). In WT, VF increased firing frequency, whereas no significant change was observed in *oxt, oxtra*, and *c1qb* mutant females. Mean ± SEM, n = 6, 6, 4, 4, 5, 5, 4, 4. Mann–Whitney U test: *P < 0.05; n.s., not significant.

### Impaired activity responses of TN-GnRH3 neurons in mutant medaka

To determine whether TN-GnRH3 neurons are functionally altered in these mutants, we measured their spontaneous firing activity using loose cell-attached patch-clamp recordings from GFP-positive TN-GnRH3 neurons (Fig. 4d). TN-GnRH3 neurons are known to exhibit a low basal pacemaker frequency and exhibit low receptivity under default conditions, whereas visual presentation of a male increases their firing frequency via an autocrine GnRH3 mechanism, leading to mate acceptance^11,13^. We therefore examined whether visual exposure to a male induces an increase in spontaneous firing frequency in each genotype. In WT, TN-GnRH3 neurons showed a significant increase in firing frequency upon male presentation. In contrast, this increase was not observed in any of the mutant lines (*oxt, oxtra,* and *c1qb*) (Figs. 4e-f). These results indicate that TN-GnRH3 neurons exhibit impaired activity responses in these mutants.

## Discussion

In this study, we demonstrate that oxytocin signaling is required for proper maturation of TN-GnRH3 circuits underlying mate preference in medaka. Disruption of oxytocin signaling resulted in impaired mate preference, accompanied by reduced expression of complement component C1q, altered microglial morphology, abnormal TN-GnRH3 projections, and impaired neuronal activity. Notably, disruption of c1qb phenocopied both behavioral and neuronal defects, supporting a functional link between oxytocin signaling and complement-mediated processes. Consistent with the role of complement signaling in synaptic refinement, our data support a model in which oxytocin signaling regulates TN-GnRH3 circuit maturation through microglia-associated synaptic mechanisms. Reduced C1q expression, together with increased innervation and smaller varicosities, is consistent with impaired maturation of neuronal projections. Microglia are key regulators of synaptic refinement, and their altered morphology in mutants further supports a disruption of this process.

Interestingly, although microglia in mutants exhibited a more ramified morphology, typically associated with homeostatic states^19^, transcriptomic reanalysis revealed reduced expression of the microglial marker p2ry12, which is commonly associated with surveillant or homeostatic-like microglial states^26^, in oxytocin signaling–deficient medaka (Fig. S5). This apparent discrepancy suggests that the ramified morphology does not necessarily reflect a typical molecular or functional state, and may indicate impaired microglial maturation or functional dysregulation.

Although our data suggest that oxytocin signaling influences microglial properties, the mechanisms underlying this altered microglial state remain unclear. Previous studies have reported oxytocin receptor expression in cultured microglia^4^; however, its expression *in viv*o remains controversial, with limited or inconsistent evidence for expression in microglia in the mouse brain^27^. Consistent with this, we did not detect oxytocin receptor expression in microglia in medaka (Fig. S2). Moreover, reductions in *c1qb* expression and alterations in microglial morphology were observed broadly across brain regions, whereas oxytocin receptor expression is spatially restricted. This spatial mismatch supports the idea that oxytocin signaling may influence microglial function indirectly, potentially via other cell types or circuit-level mechanisms. The causal relationship between reduced C1q expression and altered microglial morphology also remains unresolved. The observation that *c1qb* mutants exhibit increased microglial ramification raises the possibility that reduced complement signaling contributes to microglial alterations, although direct evidence for such a relationship remains limited. Conversely, oxytocin signaling may influence microglial developmental or activation states, which could secondarily affect C1q expression. These findings suggest that complement signaling and microglial state may be functionally coupled, and further studies will be required to determine their directionality.

Although the relationship between complement signaling and microglial state remains unclear in oxytocin signaling–deficient mutants, these alterations are likely to affect TN-GnRH3 circuit organization. TN-GnRH3 neurons form a tightly packed cluster that supports coordinated neuronal activity. Disruption of this organization in mutants is therefore likely to impair coordinated output and may contribute to the failure of TN-GnRH3 neurons to increase their firing frequency in response to visual stimuli. A notable finding of this study is the dissociation between TN-GnRH3 neuronal activity and behavioral output. While visual exposure to a male increased firing frequency in WT fish, this response was absent in mutants, despite their high behavioral receptivity. This suggests that TN-GnRH3 activity is not simply a permissive signal for mating receptivity, but may contribute to the appropriate gating of behavioral output in response to visual social cues. In oxytocin signaling–deficient mutants, abnormal circuit organization may disrupt both stimulus-evoked TN-GnRH3 activity and basal circuit output, resulting in high receptivity despite the absence of firing increases. This interpretation is consistent with altered synaptic maturation based on multiple structural features (Figs. 3–4). Previous studies have shown that TN-GnRH3 neurons project extensively to the optic tectum and modulate retino-tectal neurotransmission, thereby influencing visual information processing^21^. These findings suggest that TN-GnRH3 neurons play a direct role in shaping visually guided behaviors. In this context, the increased innervation and altered activity of TN-GnRH3 neurons observed in mutants may disrupt the processing of visual cues required for mate preference, a behavior that critically depends on visual information.

Together, our findings support a model in which oxytocin signaling links microglia-associated synaptic refinement to TN-GnRH3 circuit maturation, thereby ensuring proper coupling between neuronal activity and behavior (Fig. 5). The accessibility of TN-GnRH3 circuits in medaka provides a unique opportunity to link molecular, cellular, and circuit-level mechanisms to behavior.

**Fig. 5.**
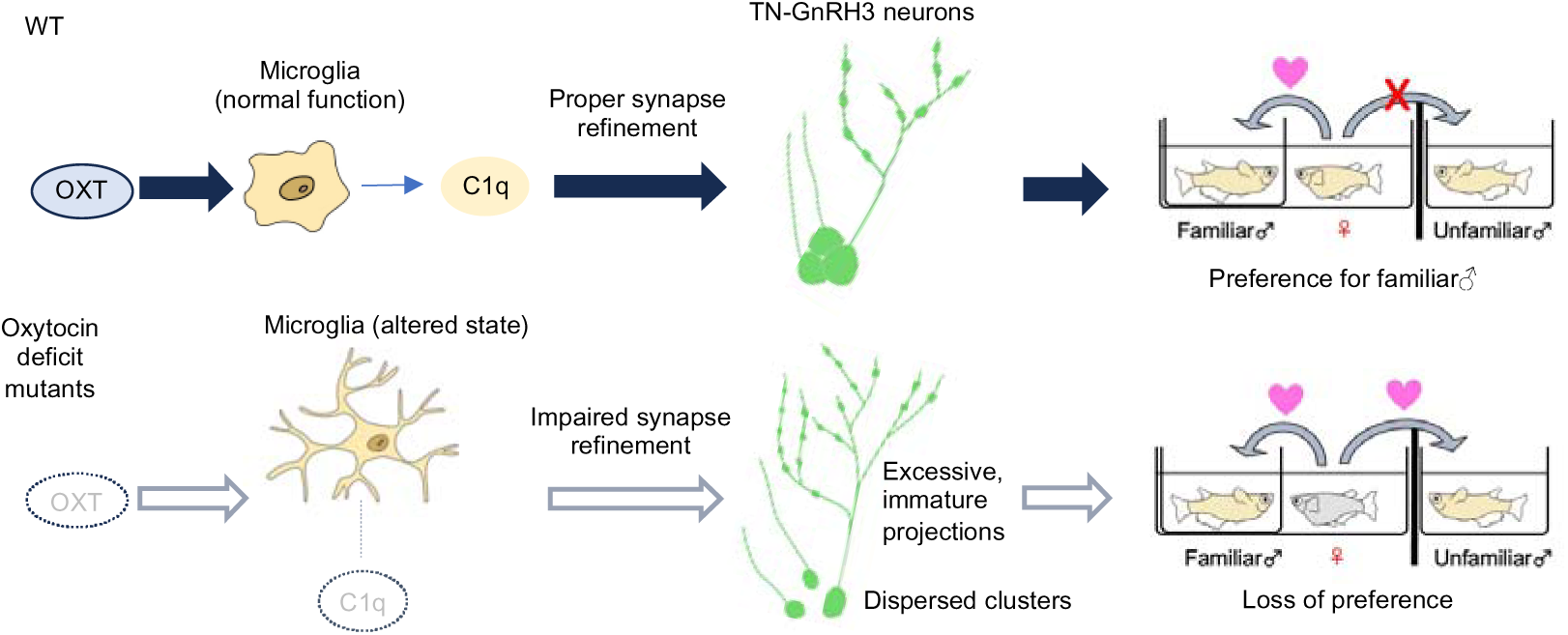
Model of oxytocin-dependent regulation of TN-GnRH3 circuits and mate preference. Model illustrating how oxytocin signaling regulates mate preference through microglia-associated synaptic mechanisms of TN-GnRH3 circuits. In WT, oxytocin signaling promotes normal microglial function and *c1q* expression, supporting proper synaptic refinement and organized TN-GnRH3 projections, leading to normal mate preference. In oxytocin signaling–deficient mutants, reduced *c1q* expression and altered microglial states are associated with impaired synaptic refinement, excessive immature projections, and dispersed TN-GnRH3 neuronal clusters, ultimately leading to loss of mate preference.

In contrast to the role of oxytocin signaling in microglia-associated synaptic processes suggested by our model, most previous studies have examined its effects on microglia under inflammatory conditions, where it suppresses microglial activation. Our findings instead suggest that oxytocin signaling may support microglial function under non-inflammatory conditions. This difference may reflect the distinct roles of oxytocin in homeostatic versus pathological contexts, which have not been extensively explored. Beyond these differences in microglial regulation, oxytocin is widely recognized for its acute neuromodulatory roles in regulating social behaviors in adults^28,29^, our findings raise the possibility that oxytocin signaling also contributes to the developmental maturation of neural circuits. Rather than acting solely as a transient modulator of neuronal activity, oxytocin may shape the structural and functional properties of circuits through microglia-associated synaptic refinement. In addition, studies in rodents have implicated complement component C1q in neurodevelopmental processes such as synaptic pruning^7,8^, as well as in the pathogenesis of neurodevelopmental disorders including schizophrenia^30^. Both excessive and insufficient synaptic refinement have been associated with abnormal circuit development^31^.

Our findings provide a framework for understanding how oxytocin signaling contributes to neural circuit maturation through microglia-associated synaptic refinement. These results reveal a previously unrecognized role for oxytocin in regulating microglial function and circuit development. Further investigation will be required to determine whether similar mechanisms operate in mammals and how their dysregulation may relate to neurodevelopmental conditions.

## Materials and methods

### Ethics statement

All procedures and experimental protocols involving live medaka fish (*Oryzias latipes*) were conducted in strict accordance with the guidelines approved by the Animal Care and Use Committee of Hokkaido University (permit no. 23-0115). To ensure the well-being of the animals, surgeries were carried out under MS-222 anesthesia, adhering to the principles outlined in the NIH Guide for the Care and Use of Laboratory Animals. Diligent efforts were taken to minimize animal discomfort and distress throughout the study.

### Fish and breeding conditions

Medaka fish (*Oryzias latipes*; OK-cab strain and mutants) were kept in groups in plastic aquariums (16 cm × 25 cm × 14 cm [height]). All fish were hatched and bred within our laboratory facilities. The oxytocin (*oxt*) and oxytocin receptor (*oxtra*) mutant medaka lines used in this study were previously generated and characterized^14^. Briefly, the *oxt* mutant carries a point mutation (I22F), and the *oxtra* mutant was generated using CRISPR/Cas9-mediated genome editing. For the experiments, sexually mature female medaka aged between 4 to 5 months were used. The water temperature was maintained at approximately 28 °C, and a consistent photoperiod was provided, with 14 hours of light from 0800 to 2200 daily, using standard fluorescent lighting.

### CRISPR/Cas9 experiment for *c1qb* mutant generation

To create *c1qb* mutants in medaka, we employed the CRISPR/Cas9 system^32^. A guide RNA (gRNA) was designed using the CRISPR/Cas9 target online predictor available at COS (https://aceofbases.cos.uni-heidelberg.de:8044/index.html). The sequence “AACGGCAAGCAAGGTCCTAAAGG” (with the underlined section representing the PAM sequence) was selected as the target. For gRNA formation, Alt-R crRNA and Alt-R tracrRNA (IDT) were combined in equimolar amounts in IDT Duplex Buffer, heated to 95°C for 5 minutes, then gradually cooled to room temperature. Alt-R S.p. HiFi Cas9 Nuclease V3 protein (250 ng/μl) and sgRNA (50 ng/ul) were mixed and injected into fertilized eggs of the cab strain by a microinjector. We backcrossed CRISPR/Cas9 mutants with cab fish twice and then crossed those fish to generate the homozygous mutants. The genotype of each fish was determined by direct sequencing. We performed polymerase chain reaction (PCR) with *c1qb*-specific primers: forward primer 5’ - CATCAGTATTATGGTAAGAGGCTTTTAC -3’ and reverse primer 5’ - GAGGTCATGCTAGACACAGGTGAG -3’.

### Female mating receptivity assay

A female mating receptivity assay was performed as previously described^13,14^. On the day before the assay, males and females were separated in the evening (20:00–21:00) under two conditions: visual familiarization and no visual familiarization. On the following morning (10:00–12:00), the divider was removed and a male was introduced into the female’s tank. Mating behavior was recorded for 5 min. Based on video recordings, the timing of male quick-circle courtship displays and subsequent wrapping behavior followed by spawning was determined. The latency to mate was defined as the interval between the first male courtship display and the wrapping event leading to spawning. Latency to mate was compared between groups, with shorter latency indicating higher female mating receptivity.

### *In situ* hybridization

The expression patterns of *c1qb* and *oxtra* were visualized using *in situ* hybridization on frozen brain sections, following previously described methods^14,33^. The antisense RNA probe for *oxtra* was identical to that used in the previous study^14^. The cDNA fragments for c1qb was amplified using synthesized cDNA, which was prepared from mRNA extracted from the medaka brain. For this, we used the ReverTra Ace qPCR RT Master Mix with gDNA Remover (TOYOBO). The primer sequences used were as follows: *c1qb* forward primer 5’ -TCCAGGGATACCCGGAACCCATG-3’, *c1qb* reverse primer 5’ -TCAGGCGGTGGGGAAGATCAG -3’. The amplified cDNA of *c1qb* was then cloned into the pCR2.1 TOPO vector (Thermo Fisher Scientific). From these constructs, digoxigenin (DIG)-labeled riboprobes were synthesized using T7 or SP6 polymerase along with a DIG labeling mix (Roche Diagnostics). For the detection of hybridization signals, two methods were employed. Alkaline phosphatase (AP) staining was performed using anti-DIG-AP antibody (diluted at 1:1000, 11093274910; Roche Diagnostics) and 5-bromo-4-chloro-3-indolyl phosphate/nitro blue tetrazolium (BCIP/NBT) substrate (Roche Diagnostics) for chromogenic detection. The total number of *c1qb*-positive cells in the midbrain and forebrain was quantified using two adjacent sections per individual. Manual counting was independently performed by two investigators, and the averaged values were used for analysis. The mean cell numbers were compared between WT and three mutant groups. For fluorescent detection, we used anti-DIG mouse monoclonal antibody [21H8] (diluted at 1:200, ab420; Abcam) as the primary antibodies and Cy3-conjugated goat anti-mouse secondary antibody (diluted at 1:200, #AP124C; Merck Millipore) for detection, with confocal microscopy (LSM900; Zeiss) utilized for imaging. The classification and naming of the brain regions in our study were consistent with those listed in the medaka brain atlases.

### Double labeling with *in situ* hybridization and immunohistochemistry

The double labeling procedure was similar to the *in situ* hybridization protocol, with the addition of immunohistochemistry steps^33^. A mixture of anti-DIG mouse monoclonal antibody [21H8] (diluted at 1:200, ab420; Abcam) and rabbit anti-Iba1 antibody (diluted at 1:200, 019-19741; Wako) served as the primary antibody, and a mixture of Cy3-conjugated goat anti-mouse secondary antibody (diluted at 1:200, #AP124C; Merck Millipore) and Alexa Fluor 488-conjugated goat anti-rabbit secondary antibody (diluted at 1:250, A-11008; Invitrogen) as the secondary antibody.

### Generation of transgenic medaka using Tol2 transposition

A Tol2-based gnrh3:GFP expression construct was generated using the destination vector pDestTol2pA2 from the Tol2kit^34^. The gnrh3 promoter fragment^16^ was amplified from medaka genomic DNA, while the vector backbone and GFP fragment were amplified from pDestTol2pA2 and pME-EGFP (Tol2kit), respectively. These fragments were assembled into the Tol2 backbone using NEBuilder HiFi DNA Assembly (New England Biolabs).

Primers used for amplification contained homologous overhangs for HiFi assembly. The full primer sequences are as follows (5′–3′): gnrh3 promoter F, ctgaaacacaggccagatgggCAAGGTCGGTTTGAAGGTCGGAG; gnrh3 promoter R, tcgcccttgctcaccatggtgTCTGTGTTTTCTCTGTTCGCATG; vector F, ctggccgtcgttttacTCTGCGAAGATACGGCCACG; vector R, ctccgaccttcaaaccgaccttgCCCATCTGGCCTGTGTTTCAG; GFP F, catgcgaacagagaaaacacagaCACCATGGTGAGCAAGGGCGA; GFP R, cgtggccgtatcttcgcagaGTAAAACGACGGCCAG.

Lowercase sequences indicate homologous overlaps used for HiFi DNA Assembly, and uppercase sequences correspond to gene-specific regions.

The Tol2 construct (10 ng/µl) and Tol2 transposase mRNA (10 ng/µl) were co-injected into one-cell stage embryos. Approximately 1–2 nL of the injection solution was delivered into the embryos. Tol2 transposase-encoding capped mRNA was synthesized by in vitro transcription using the SP6 mMessage mMachine Kit (Ambion, AM1340) from pCS2+Tol2 plasmid linearized with NotI. The synthesized RNA was purified using the RNeasy kit (Qiagen). GFP expression was screened under a fluorescence stereomicroscope. Injected embryos were raised to adulthood and screened for germline transmission. Stable transgenic lines were established by outcrossing to WT fish and subsequently crossed with mutant fish for analysis.

### Immunohistochemistry with cryosections

Cryosections were permeabilized in cold methanol (−20°C) for 5 min and washed in TBST. Sections were blocked using blocking reagent (Roche) in TBST for 5–10 min at room temperature. Sections were incubated with primary antibody (goat anti-Iba1, 1:200, NB100-1028; Novus or rabbit anti-GFP, 1:200, 598; MBL) for 1 h at room temperature, followed by washing in TBST (3 × 5 min). Sections were then incubated with fluorescent secondary antibody (donkey anti-goat Cy3, 1:250, AP180C; Sigma-Aldrich or goat anti-rabbit Alexa Fluor 488, 1:250, A-11008; Invitrogen) and DAPI (1:1000) for 30 min at room temperature in the dark. After washing (3 × 5 min in TBST), sections were mounted with PVA mounting medium. Fluorescence images at low magnification were acquired using a fluorescence stereomicroscope, whereas high-magnification images were obtained using a confocal microscope. The total number of Iba1-positive cells in the midbrain and forebrain was quantified using two adjacent sections per individual. Manual counting was independently performed by two investigators, and the averaged values were used for analysis. The average microglial cell area was calculated by measuring the total Iba1-positive area using Fiji (ImageJ) and dividing it by the number of Iba1-positive cells counted in the same sections.

The proportion of GFP-positive area within the OT or Dl was quantified using Fiji (ImageJ). Regions of interest (ROIs) corresponding to the OT or Dl were manually delineated, and the area of GFP-positive signal within each ROI was measured. The percentage of GFP-positive area was calculated as the ratio of GFP-positive area to the total ROI area. Quantification was performed using two adjacent sections per individual, and the averaged values were used for analysis.

Varicosity size and density were quantified from axonal segments in the OT or Dl. For each individual, five axonal segments longer than 50 μm were randomly selected, and varicosities were identified using the Analyze Particles function in Fiji (ImageJ). The mean varicosity size and the number of varicosities per μm (density) were calculated for each segment, and the averaged values across the five segments were used for analysis.

### Immunohistochemistry with vibratome sections

Adult medaka brains from gnrh3:GFP transgenic fish were fixed by perfusion with 4% paraformaldehyde (PFA), dissected, and post-fixed overnight at 4°C. Samples were washed in PBS and stored in methanol at − 20°C. For sectioning, samples were rehydrated through a graded methanol series into PBS and embedded in 3% agarose. Embedded brains were sectioned at 100 μm thickness using a vibratome, and sections containing GFP-positive signals were selected for analysis. Sections were rehydrated and blocked using blocking reagent (Roche) in TBST for 1 h at room temperature, followed by incubation with primary antibody (rabbit anti-GFP, 1:750, 598; MBL) overnight at 4°C. After extensive washing, sections were incubated with fluorescent secondary antibody (anti-rabbit Alexa Fluor 488, 1:500) and DAPI (1:1000) for 4–6 h. After washing, sections were post-fixed in 4% PFA, washed, and mounted in glycerol-based mounting medium containing DABCO. Images were acquired with confocal microscopy (LSM900; Zeiss). Sagittal sections were analyzed for gnrh3 neuronal cell body morphology. For each individual, only sections in which all gnrh3 neuronal cell bodies (a bilateral pair) were contained within a single section were selected for analysis, to enable accurate quantification of cluster organization. The distance between the farthest cell bodies (μm) and the nuclear area (μm²) were measured using ZEN software (Zeiss). Nuclear area was measured for four randomly selected cells per individual and averaged to obtain a representative value for each individual.

### Electrophysiology

Loose cell-attached patch-clamp recordings were performed as previously described^13^ with minor modifications. Adult medaka were anesthetized by immersion in 0.02% MS-222 and decapitated. Brains were immediately isolated in MS-222-free extracellular solution and maintained in this solution throughout subsequent preparation, including removal of the ependymal layer of the telencephalon to expose GnRH3-GFP-positive neurons and mounting in the recording chamber. Recordings were started at least 20 min after transfer to MS-222-free solution and were performed at room temperature. Recordings were performed under an upright fluorescence microscope equipped with infrared differential interference contrast optics, allowing visual identification of GFP-positive neurons. Electrical activity was monitored from GFP-positive neurons using a loose-patch pipette (resistance, ∼3 MΩ). Patch pipettes were pulled from borosilicate glass capillaries (1.5 mm outer diameter; Narishige) using a micropipette puller (P-1000; Sutter Instruments). The solution for the loose-patch pipette contained 150 mM NaCl, 3.5 mM KCl, 2.5 mM CaCl_2_, 1.3 mM MgCl_2_, 10 mM HEPES, and 10 mM glucose (pH 7.4, adjusted with NaOH). A low-resistance seal (<100 MΩ) was formed after gentle release of positive pressure. Loose cell-attached recordings were obtained using a MultiClamp 700B amplifier (Molecular Devices) in I = 0 mode, without current injection.

Firing frequency was analyzed using AxoGraph software. Pacemaker activity was quantified by calculating the mean firing frequency over a 5 min period starting 1 min after the onset of recording. Firings were detected and counted automatically.

### Reanalysis of RNA-seq data

Previously generated RNA-seq data from WT, *oxt*, and *oxtra* mutant female medaka brains^14^ were reanalyzed to assess expression levels of *p2ry12*. Read coverage across the *p2ry12* locus was visualized using IGV.

### Statistical analysis

Behavioral indices in the female mating preference assay were compared using the Mann–Whitney U test implemented in R. For histological analyses, comparisons between WT and mutant groups were performed using the Kruskal–Wallis test followed by Steel’s post hoc test implemented in R. In patch-clamp experiments, pacemaker frequency was compared between visual familiarization conditions using the Mann–Whitney U test implemented in R. All statistical tests were two-tailed, and p values less than 0.05 were considered statistically significant.

## Acknowledgements

We thank K. Yamanaka for kindly providing the anti-Iba-1 rabbit antibody; T. Deguchi, T. Tani and Y. Kamei for constructive discussions; the BioResource Project Medaka (https://shigen.nig.ac.jp/medaka) for providing the OK-cab strain (strain ID:MT830). This work was supported by KAKENHI Grant Numbers 21H05708 (SY), 23K05841 (SY), 23H03839 (SY), JST FOREST Program (JPMJFR241Y) (SY), Astellas Foundation for Research on Metabolic Disorders (SY), Takeda Science Foundation (SY) and the Naito Foundation (SY).

## Author contributions

K. N., H. T., S. N. and S.Y. designed research; M. H., M. O. and S.Y. performed research; K. H., F. W., S. H., T. A. and M. M. contributed to electrophysiological experiments; M. H., M. O. and S.Y. analyzed data; S.Y. wrote the paper. All authors reviewed the manuscript.

