## Supplemental Figures for "Oxytocin regulates TN-GnRH3 circuit maturation and mate preference through C1q-dependent synaptic mechanisms"

**Supplementary figure legend**

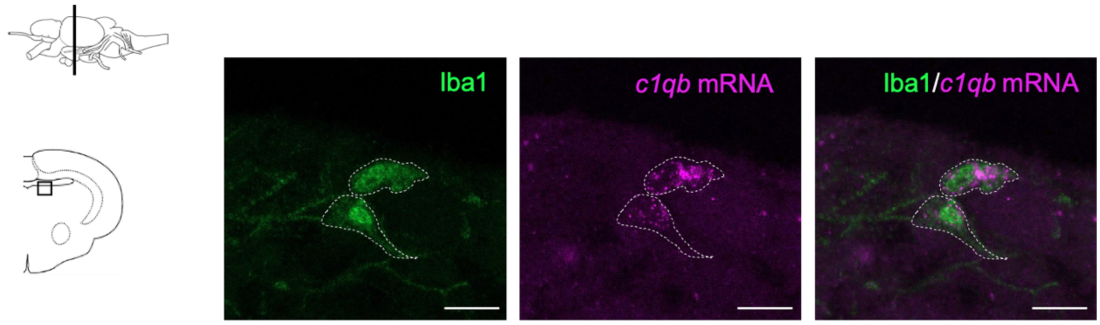


**Fig. S1 *c1qb* expression in microglia in medaka brain**

Representative images of combined immunostaining for Iba1 (green) and *in situ* hybridization for *c1qb* mRNA (magenta) in the medaka brain. Merged images show *c1qb* signals in Iba1-positive microglial cells. Dashed outlines indicate representative microglial cells. The schematic on the left shows the coronal section level and the analyzed region. Scale bars represent 10 µm.

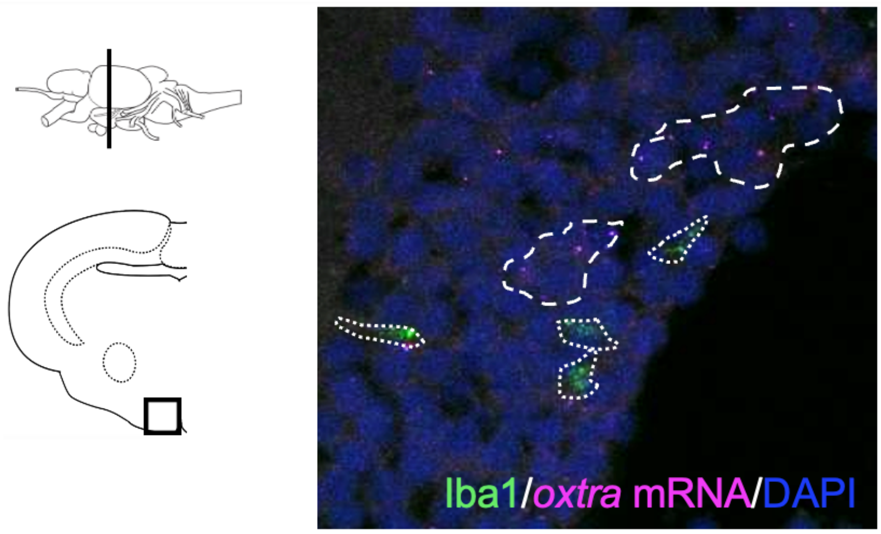


**Fig. S2. *oxtra* expression is not detected in microglia in the medaka brain**

Representative images of combined immunostaining for Iba1 (green) and *in situ* hybridization for *oxtra* mRNA (magenta) with DAPI (blue). *oxtra* mRNA signals were observed in cells distinct from Iba1-positive microglia. Dashed outlines indicate representative *oxtra*-positive cells, and dotted outlines indicate Iba1-positive microglia. The analyzed region corresponds to a brain area where *oxtra* expression has been previously reported[14]. Left, schematic of the coronal section level and analyzed region.

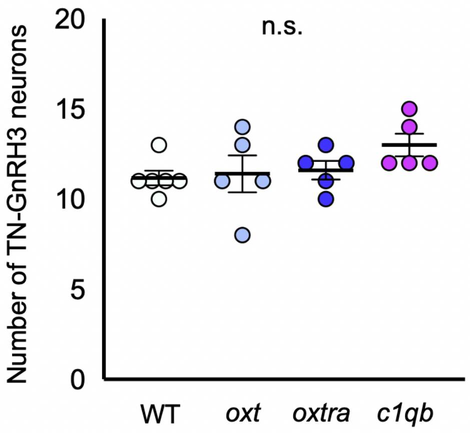


**Fig. S3. TN-GnRH3 neuronal soma number is unchanged in mutant medaka**
Quantification of the number of TN-GnRH3 neuronal somata, in WT, *oxt* mutant, *oxtra* mutant, and *c1qb* mutant females. Mean ± SEM, n = 6, 5, 5, 5. Kruskal–Wallis test. n.s., not significant.

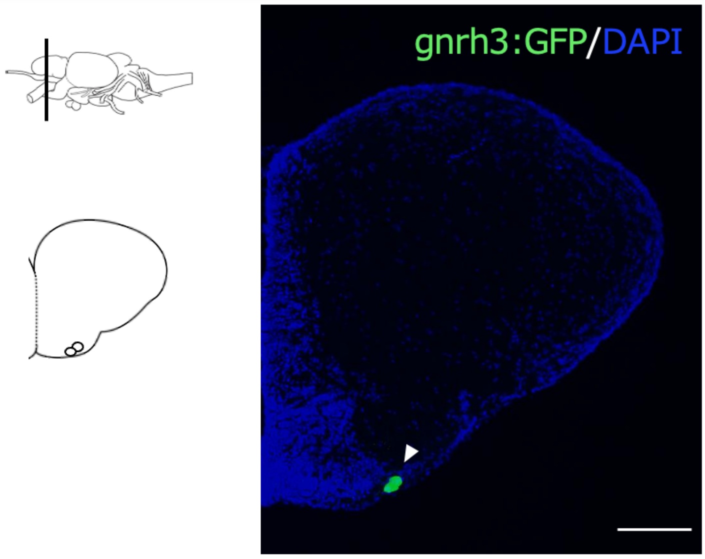


**Fig. S4. Coronal section level containing TN-GnRH3 neuronal somata.**
Representative image of gnrh3:GFP (green) with DAPI (blue) in the medaka brain. Arrowhead indicates TN-GnRH3 neuronal somata. Schematics on the left show the coronal section level and the position of the analyzed region. The scale bar represents 10 μm.

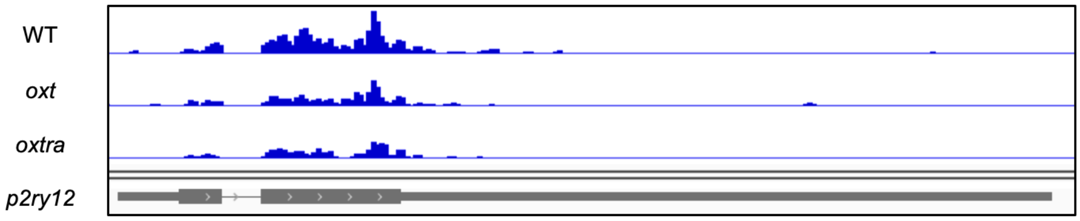


**Fig. S5. Reduced expression of *p2ry12* in oxytocin signaling–deficient medaka.** Genome browser view showing RNA-seq read coverage across the *p2ry12* locus in WT, *oxt* mutant, and *oxtra* mutant females. Reduced read coverage is observed in *oxt* and *oxtra* mutants compared to WT.
